# Association between proteomic blood biomarkers and DTI/NODDI metrics in adolescent football players

**DOI:** 10.1101/2020.02.20.958694

**Authors:** Keisuke Kawata, Jesse A. Steinfeldt, Megan E. Huibregtse, Madeleine K. Nowak, Jonathan T. Macy, Andrea Shin, Zhongxue Chen, Keisuke Ejima, Kyle Kercher, Sharlene D. Newman, Hu Cheng

## Abstract

The objective of the study was to examine the association between diffusion MRI techniques [diffusion tensor imaging (DTI) and neurite orientation/dispersion density imaging (NODDI)] and brain-injury blood biomarker levels [Tau, neurofilament-light (NfL), glial-fibrillary-acidic-protein (GFAP)] in high-school football and cross-country runners at their baseline, aiming to detect cumulative neuronal damage from prior seasons. Twenty-five football players and 8 cross-country runners underwent MRI and blood biomarker measures during preseason data collection. The whole-brain, tract-based spatial statistics was conducted for six diffusion metrics: fractional anisotropy (FA), mean diffusivity (MD), axial/radial diffusivity (AD, RD), neurite density index (NDI), and orientation dispersion index (ODI). Diffusion metrics and blood biomarker levels were compared between groups and associated within each group. The football group showed lower AD and MD than the cross-country group in various axonal tracts of the right hemisphere. Elevated ODI was observed in the football group in the right hemisphere of the corticospinal tract. Blood biomarker levels were consistent between groups except for elevated Tau levels in the cross-country group. Tau level was positively associated with MD and negatively associated with NDI in the corpus callosum of football players, but not in cross-country runners. Our data suggest that football players may develop axonal microstructural abnormality. Levels of MD and NDI in the corpus callosum were associated with serum Tau levels, highlighting the vulnerability of the corpus callosum against cumulative head impacts. Despite observing multimodal associations in some brain areas, neuroimaging and blood biomarkers may not strongly correlate to reflect the severity of brain damage.

## INTRODUCTION

Concussive and subconcussive brain injury in sports have emerged as a complex public health issue. Policy and rule changes, as well as societal awareness, have played a catalytic role in decreasing concussion incidence in sports.^1^ However, despite decades of investigation, there is no concrete evidence on gold-standard diagnostic biomarkers for concussion, preventive tools that can increase neural resiliency to trauma, or factors contributing to the potential long-term consequence of subconcussive head impact exposure. This knowledge gap is partly due to the unimodal approach, in which many papers report data derived from a single modality (e.g., neuroimaging, blood biomarker, behavioral measures). This precludes validation of study findings. For example, elevated Tau protein in blood theoretically indicates axonal damage or degeneration, but without cross-referencing against imaging data, the usefulness of tau protein as a surrogate for brain damage remains speculative at best.

Several interdisciplinary groups have begun testing 2- and 3-way multimodal relationships that reflect subconcussive neuronal stress. In 2014, initial studies by Talavage et al.^2^ and Bazarian et al.^3^ revealed head impact-dependent declines in neural activation patterns and axonal microstructural integrity after a single high school and college football season, respectively. These neuroimaging findings were correlated with declined cognitive function^2, 4^ and the development of autoimmune response to brain-derived blood biomarkers (e.g., ApoA1, S100B).^3, 5^ Despite the unequivocal importance of the multimodal approach, association studies in subconcussion research are limited.^6, 7^

Neuroimaging techniques, especially diffusion MRI, and brain-derived blood biomarkers are the fastest growing areas of neurotrauma research. Diffusion tensor imaging (DTI) is the most extensively used technique worldwide to examine the white matter microstructural properties in humans. However, DTI metrics such as mean diffusivity (MD) and fractional anisotropy (FA) represent basic statistical descriptions of diffusion that do not directly correspond to biophysical properties of neuronal axons.^8^ In 2012, Zhang et al.^9^ introduced the neurite orientation and dispersion density imaging (NODDI) technique that can measure axonal density within white matter, dispersion of axonal orientation, and free water diffusion. The combined use of DTI and NODDI has been shown to detect progressive axonal degeneration even 6 months after a concussion.^10^ Similar to neuroimaging techniques, blood biomarker technology has evolved to be able to detect neural factors at a femtomolar concentration. Among the many potential biomarkers for brain injury, Tau, neurofilament-light (NfL), and glial fibrillary acidic protein (GFAP) have shown their superior ability to predict concussion recovery time,^11, 12^ cumulative subconcussive axonal damage,^13–15^ and absence of intracranial bleeding.^16–18^ However, the relationships between DTI/NODDI metrics and blood biomarkers in reflecting cumulative neural stress from football head impacts have never been reported in the literature.

Therefore, we conducted a cross-sectional association study in high school football players and cross-country runners (control athletes) to examine the relationship between axonal diffusion imaging metrics and blood biomarkers at their preseason baseline, aiming to detect cumulative neuronal damage from prior football seasons. We hypothesized that there would be group differences in diffusion metrics and blood biomarkers. We also hypothesized that there would be significant associations between imaging and blood biomarkers to reflect axonal microstructural damage in some areas of the brains of football players, but not in cross-country runners.

## METHODS

### Participants

This single-site, cross-sectional study enrolled 25 male high school football athletes and 8 male high school cross-country athletes, who served as the control group. None of the 33 participants was diagnosed with a concussion or traumatic brain injury in the 12 months prior to the enrollment. Inclusion criterion was being an active high school football or cross-country team member. Exclusion criteria included a history of head and neck injury in the previous year or neurological disorders. Conditional exclusion criteria for the neuroimaging data collection were metal implants in the body or implanted electro/magnetic devices (e.g. orthodontic braces, pacemakers, aneurysm clips). The Indiana University Institutional Review Board approved the study, and all participants and their legal guardians gave written informed consent. The data were collected during the preseason baseline assessment in July 2019 and included self-reported demographic information (age, race/ethnicity, height, weight, number of previously diagnosed concussions, and years of tackle American football experience), 7 mL of blood samples, and MRI scans.

### Blood biomarker assessments

Seven-milliliter samples of venous blood were collected into red-cap serum vacutainer sterile tubes (BD Bioscience). Blood samples were allowed to clot at room temperature for a minimum of 30 min. Serum was separated by centrifugation (1,500 x g, 15 min) and stored at - 80°C until analysis. Serum levels of Tau, NfL, and GFAP were measured using the Simoa^TM^ Platform (Quanterix), a magnetic bead-based, digital enzyme-linked immunosorbent assay (ELISA) that allows detection of proteins at femtomolar concentrations.^19^ An analytical protocol was previously described in detail.^20^ The analyses were performed by a board-certified laboratory technician blinded to the study design and subject characteristics. Limit of detection was 0.024 pg/mL for Tau, 0.104 pg/mL for NfL, and 0.221 pg/mL for GFAP. The average intra-assay coefficients of variation for the samples were 6.7 ± 5.2% for Tau, 8.3 ± 6.0% for NF-L, and 3.7 ± 2.7% for GFAP.

### MRI acquisition

The MRI data were acquired on a 3T Siemens Prisma MRI scanner (Siemens, Erlangen, Germany) equipped with a 64-channel head/neck coil. High-resolution anatomical images (T1 weighted) were acquired using 3D MPRAGE pulse sequence with the following parameters: TR/TE=2400/2.3 ms, TI=1060 ms, flip angle=8, matrix=320×320, bandwidth=210 Hz/pixel, iPAT=2, resulting in 0.8 mm isotropic resolution. For diffusion analysis, two consecutive diffusion weighted imaging (DWI) sessions with opposite phase encoding directions were performed with a simultaneous multi-slice single-shot spin-echo echo-planar pulse sequence with the following parameters: TE=89.4 ms; TR=3590 s, flip angle=90, 1.5 mm isotropic resolution. Each session had 103 images with different diffusion weightings and gradient directions summarized as following: 7 b=0 s/mm^2^, 6 directions with b=500 s/mm^2^, 15 directions with b=1000 s/mm^2^, 15 directions with b=2000 s/mm^2^, and 60 directions b=3000 s/mm^2^.

### Imaging processing

First, the DWI images were denoised using the PCA-based denoising tool in Mrtrix (https://www.mrtrix.org/),^21^ and then magnetic field map information for susceptibility artifacts correction was derived from the b0 (b=0 s/mm^2^) images with opposite phase encoding directions using TOPUP in FSL (https://fsl.fmrib.ox.ac.uk/fsl/fslwiki).^22^ The images were then corrected for susceptibility artifact, eddy current distortions, and motion artifacts simultaneously using the “eddy” command of FSL and the average of the b0 volumes as a reference. DTI analysis was performed using the FSL Diffusion Toolbox. The diffusion metrics of fractional anisotropy (FA), mean diffusivity (MD), axial diffusivity (AD), and radial diffusivity (RD) maps were calculated.

Meanwhile, the NODDI metrics including neurite density index (NDI) and orientation dispersion index (ODI) were derived using the NODDI Matlab toolbox v1.01 (http://mig.cs.ucl.ac.uk/index.php?n=Tutorial.NODDImatlab) using the default settings. NDI primarily represents axonal density within white matter, and ODI represents organization of white matter tracts.^10^

### Statistical analysis

The whole-brain, tract-based spatial statistics (TBSS) was conducted for six diffusion metrics: FA, MD, AD, RD, NDI, and ODI in FSL.^23^ The FA maps were co-registered to a template via nonlinear transformation. A skeleton of mean white matter tracts was obtained, and FA values of nearby voxels were projected to the template to obtain skeletonized FA maps. The nonlinear warps and skeleton projection derived from FA maps were applied to all other diffusion metrics to obtain skeletonized maps of MD, AD, RD, NDI, and ODI as well.

Two different statistical analyses were conducted to examine group differences between football players and cross-country runners, as well as the relationships between diffusion metrics and blood biomarkers. First, group difference in blood biomarker levels was tested by Welch’s t-test, while two-sample t-tests were used for each diffusion metric through randomized permutation.

The second analysis used complete samples to test the relationship between blood biomarker and MRI data. Univariate regression analyses were conducted for each diffusion metric in both groups against their blood biomarker levels via randomized permutation. The model included years of tackle football experience and number of concussion occurrences as covariates. The Threshold-Free Cluster Enhancement (TFCE) option was used in the permutation test, which gives cluster-based thresholding for family wise error (FEW) correction.^24^ As a result, the TFCE p-value images obtained were fully corrected for multiple comparisons across space. When there was a significant association, post-hoc analysis using a Pearson correlation coefficient was computed between the blood biomarker level and an average value of the imaging voxels that showed a significant effect in the regression analysis.

## RESULTS

### Demographics

Five of 25 participants in the football group were excluded from MRI due to a metal implant in the body (n=4) and orthodontic braces (n=1), whereas all 8 cross-country runners completed the MRI. Three serum samples in the football group were not assessed for biomarkers due to hemolysis, and a boxplot analysis identified 1 data point for Tau and NfL in the cross-country group to be an unexplainable outlier, which was excluded from the analysis. As a result, 28 imaging data (n=20 football, n=8 cross-country) and 30 blood biomarker data (n=22 football, n=7 cross-country) were valid for the group difference analysis. We used 24 completed sets of the imaging-blood biomarker data (n=17 football, n=7 cross-country) for the association analysis. Demographic information is detailed in Table 1.

**Table 1.** Group demographics

| **Table 1. Group demographics** |  |  |
| --- | --- | --- |
| **Variables** | **Football** | **Cross-Country** |
| n | 25 | 8 |
| Sex (%) | 25M (100) | 8M (100) |
| Age, y | 15.8 ± 1.11 | 15.3 ± 1.4 |
| BMI, kg/m<sup>2</sup> | 27.4 ± 6.08 | 19.3 ± 1.02 |
| No. of previous concussion |  |  |
| 0, n (%) | 17 (68.0) | 0 (87.5) |
| 1, n (%) | 6 (24.0) | 1 (12.5) |
| 2, n (%) | 2 (8.0) | 0 (0) |
| Tackle football experience, y | 5.8 ± 2.9 | 0.0 ± 0.0 |
| Race, n (%) |  |  |
| White | 21 (84) | 7 (87.5) |
| Black/African American | 0 (0) | 0 (0) |
| Asian | 0 (0) | 1 (12.5) |
| African Indiana/Alaska | 1 (4) | 0 (0) |
| Multiracial | 3 (12) | 0 (0) |
| Ethnicity, n (%) |  |  |
| Not Latino/Hispanic | 21 (84) | 6 (75) |
| Latino/Hispanic | 4 (16) | 2 (25) |
| Psychiatric condition |  |  |
| ADHD | 0 (0) | 1 (12.5)<sup>a</sup> |
| Learning Disability | 0 (0) | 0 (0) |
| Major Depressive Disorder | 0 (0) | 0 (0) |
| Blood Biomarker Levels,<br>mean±SD, pg/mL |  |  |
| Tau | 0.23 ± 0.12 | 0.45 ± 0.16 |
| Neurofilament light | 4.16 ± 1.75 | 5.00 ± 1.45 |
| Glial fibrillary acidic protein | 61.02 ± 19.59 | 76.00 ± 27.58 |
| Note: BMI, body mass index. ADHD, attention-deficit/hyperactivity disorder. <sup>a</sup> this subjects did not take medication during the baseline testing, but his MRI and biomarker data were consistent with others. |  |  |

### Group differences in diffusion metrics and blood biomarker levels

Examples of the six different diffusion metrics are shown in Fig. 1A on one slice of a representative subject that is mapped on FMRIB58_FA standard space. These parameter maps are distinct from one another, with each metric (FA, MD, AD, RD, NDI, and ODI) characterizing different diffusion features arising from underlying tissue microstructure.

**Figure 1.**
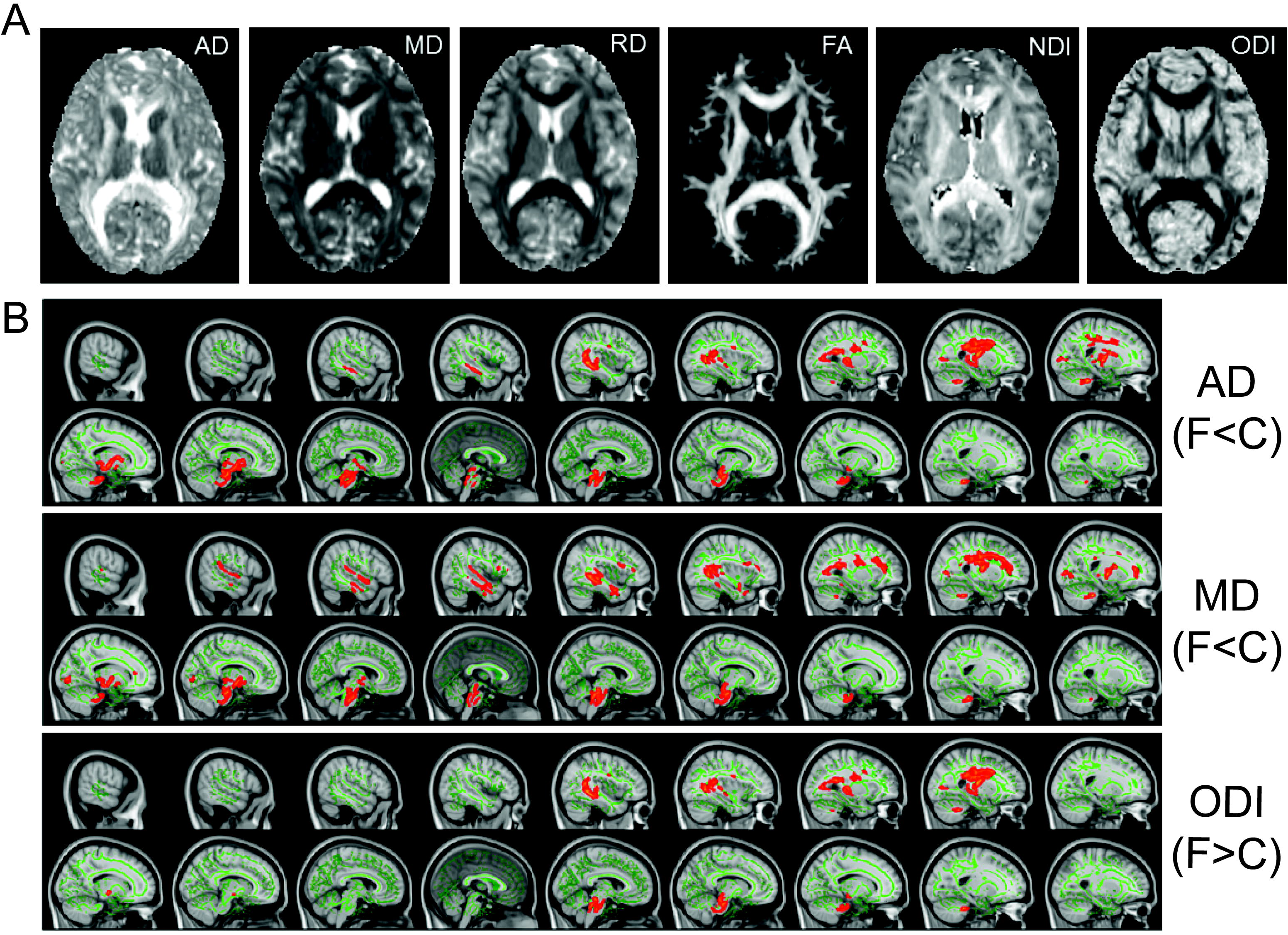
Group difference in DTI and NODDI metrics. (A) Example maps of DTI [axial diffusivity (AD), mean diffusivity (MD), radial diffusivity (RD), fractional anisotropy (FA)] and NODDI [neurite density index (NDI), and neurite orientation dispersion index (ODI)] from a single subject in FMRIB58_FA template. (B) Two-sample t-tests between football players and cross-country-runners from TBSS analysis show significant difference for AD, MD, and ODI. F, football; and C, cross-country. All results corrected for multiple comparisons using threshold-free cluster enhancement (TFCE) at p≤ 0.05.

Significant group differences were observed between football players and cross-country runners for AD, MD, and ODI (Fig. 1B). Specifically, the football group showed lower AD and MD than those of the cross-country group primarily in the right hemisphere of the inferior longitudinal fasciculus, superior longitudinal fasciculus, uncinated fasciculus, inferior fronto-occipital fasciculus, and corticospinal tract, with the lowest p-value of 0.008 for AD and 0.023 for MD. Elevated ODI was observed in the football group compared to the cross-country group, mainly in the right hemisphere of the corticospinal tract (p=0.035).

There was large variability in blood biomarker levels. The football group (0.23±0.12 pg/mL) had a significantly lower Tau level than the cross-country group (0.45±0.16 pg/mL, p=0.013: Fig 2A). There was no group difference in NfL (Football, 4.16±1.75 pg/mL vs. Cross-country, 5.00±1.45 pg/mL, p=0.219: Fig 2B) or in GFAP (Football, 61.02±19.59 pg/mL vs. Cross-country, 76.00±27.58 pg/mL, p=0.263: Fig 2C).

**Figure 2.**
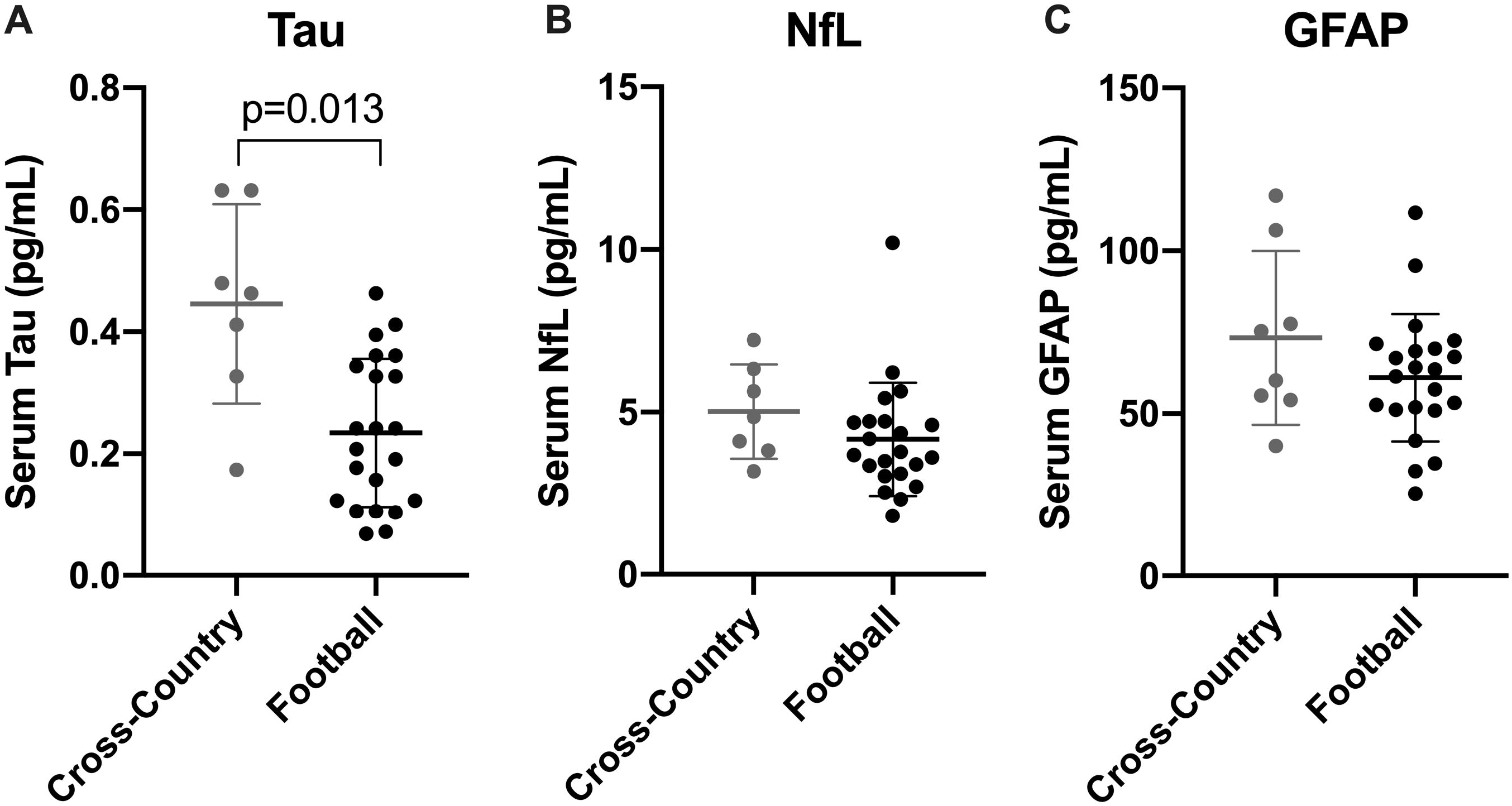
Baseline Blood biomarker levels between groups. Group difference was observed only in Tau (A), but not in NfL and GFAP.

### Associations between diffusion metrics and blood biomarker levels

Regression analyses revealed significant associations between several diffusion metrics and blood biomarker levels in the football group. Specifically, serum Tau level was positively associated with MD (Fig. 3A) and negatively associated with NDI (Fig. 3C) mainly in the corpus callosum. While the Tau-MD association was widespread over the corpus callosum (p=0.027), the Tau-NDI association was focal on the anterior body of the corpus callosum (p=0.048). In our post-hoc analysis using a Pearson correlation coefficient, we found a significant positive correlation between an average value of MD voxels that showed significant associations in Fig 3A and serum Tau levels (r=0.87, p<0.0001: Fig 3B). Similarly, there was a significant negative correlation between an average value of NDI voxels that showed significant associations in Fig 3C and serum Tau levels (r=-0.76, p=0.0005: Fig 3D). We did not observe similar relationships in the cross-country group. It is worth noting that both covariates, years of tackle football experience and number of previous concussions, had a non-significant influence on the imaging-blood biomarker association.

**Figure 3.**
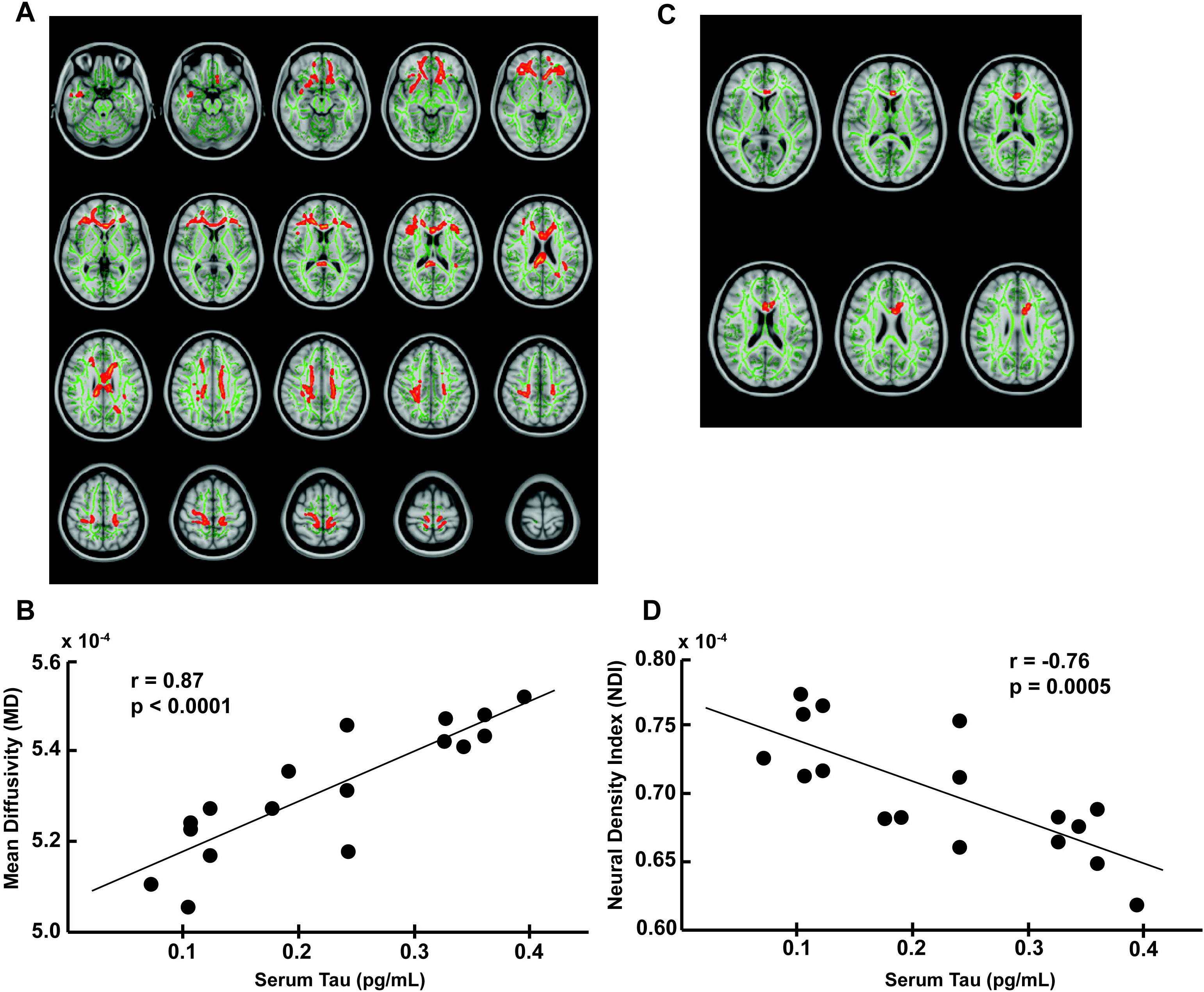
The relationship between imaging and serum Tau levels. Regression analysis in the football group showed that Tau was positively associated with mean diffusivity (A) and negatively associated with neurite density index (C) in some brain regions. Post-hoc correlation analysis revealed a positive correlation between Tau and MD that showed a significant effect in the regression analysis and a negative correlation between Tau and NDI that showed a significant effect in the regression analysis. The TFCE p-value was set to p≤0.05.

Additionally, in the football group, a small number of voxels showed positive association between GFAP and AD in the brain stem (p=0.052: Fig. 4A) and negative association between NfL and ODI in the left superior corona radiata (p=0.046: Fig. 4B).

**Figure 4.**
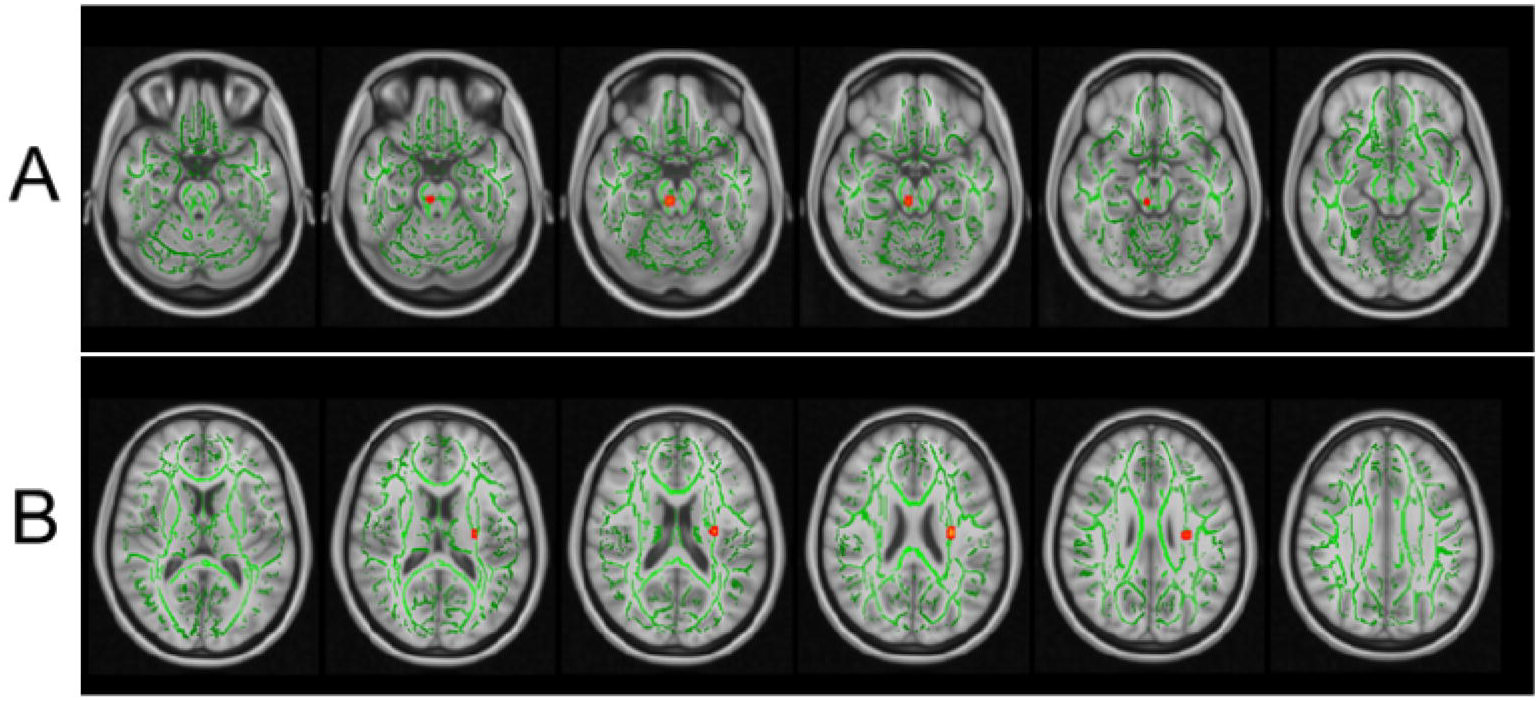
The relationship between imaging and serum GFAP levels. Regression analysis showed that GFAP was positively associated with radial diffusivity (A) and negatively associated with neurite orientation dispersion index (B) in a small portion of the longitudinal fasciculus. The TFCE p-value was set to p≤0.05.

## DISCUSSION

The novelty of the current study was the multimodal association of the sensitive neurologic metrics (DTI, NODDI, Tau, NfL, and GFAP) to reflect the potential cumulative stress in the brains of high school football players. There are three primary findings from the study: (1) Axonal microstructural integrity differed between football and cross-country runners at baseline, with a notable reduction in AD and elevation in ODI in various axonal tracts of football players. In other words, football players had less water movement along axonal tracts and disorganized axonal orientation compared to cross-country runners; (2) Football players expressed lower Tau levels in blood compared to cross-country runners; and (3) Elevated Tau levels were related to increased axonal diffusion and decreased neurite density in the corpus callosum of football players, but there was no imaging/blood biomarker relationship in cross-country runners.

The lower MD in football players, as compared to cross-country runners, is likely driven by a significantly reduced level of AD. AD represents the magnitude of diffusion along the direction of axons, and an AD reduction is often interpreted as a reflection of compromised axonal membrane integrity or axonal injury.^25, 26^ Although there are some opposing data,^27, 28^ studies have shown that contact-sport athletes had significantly lower MD in the corpus callosum than noncontact-sport athletes at preseason baseline.^29^ Previous studies further reported that significant preseason to postseason decreases in MD and AD were observed in widespread white matter areas in high school football players,^30–32^ and that the similar reduction in AD was found in youth football players at postseason.^30^ Our DTI findings were further substantiated by NODDI metrics especially in ODI, whereby football players had a significantly elevated ODI in the corticospinal tracts as compared to cross-country runners. Since the ODI reflects the overall coherence of the fibers, with lower ODI representing highly coherent structures,^9^ our data possibly indicate that playing American football may relate to chronic microstructural abnormality in axonal tracts that are important for motor control. However, it is important to note that we found no significant group differences in RD in any areas of the brain, which means that there was no evidence of demyelination in this sample of high school football players.

The corpus callosum is comprised of nearly 200 million myelinated axonal tracts that enable interhemispheric neuronal communications.^33^ The corpus callosum has been shown to be one of the most vulnerable areas of the brain to concussive and subconcussive mechanical forces (e.g., shear, stretch, shortening),^34–36^ and significant atrophy has been found in brains with chronic traumatic encephalopathy (CTE).^37, 38^ Our data on the imaging-blood biomarker associations in the corpus callosum are intriguing, in that lower NDI was related to elevated serum Tau levels in football players but not in cross-country runners. It is possible that football-related head impacts can trigger microstructural disruption in axons, as represented in reduced NDI, and concurrently induce Tau dissociation from microtubules. Dissociated Tau can reach peripheral circuitry through either blood-brain barrier leakage or glymphatic pathway.^39^ This interpretation is physiologically reasonable, except for the fact that the cross-country group had higher Tau levels than the football group.

Tau has long been regarded as a promising blood biomarker to gauge the severity of axonal damage and neurodegenerative progression because of its location and role in stabilizing microtubules. Ample clinical studies support its use for diagnosis of Alzheimer’s disease,^40^ prediction of concussion recovery duration,^11, 12^ and association with short-^41^ and long-term subconcussive neural stress.^42^ However, Tau is expressed not only in the cerebral tissue, but also in skeletal muscle and in the kidney/bladder.^43^ Di Battista et al.^44^ shed light on the contribution of the extracranial sources, such that acute high-intensity interval training could transiently elevate plasma Tau levels. Kawata et al.^45^ further corroborated the finding in football players, whereby plasma Tau levels increased after intense exercise during summer training, which possibly masked acute subconcussive effects on circulating Tau levels. Taken together, elevated Tau levels in the cross-country group may be driven by extracranial sources (e.g., muscle) from their off-season running without axonal injury, hence an NDI/Tau relationship in the corpus callosum was absent in the cross-country group.

We observed a positive association between MD and Tau in the corpus callosum of the football group. This association is noteworthy since increased MD often attributes to more severe form of TBI, whereas decreased MD is frequently reported due to repetitive subconcussive head impacts, as revealed in a recent systematic review.^46^ The corpus callosum contains less coherent axons, and it is plausible that playing football (or recurring head impacts) result in changes in MD values within the corpus callosum. Further investigation is warranted to confirm whether high levels of MD, particularly in the corpus callosum, is reflective of axonal damage, and if so to what level of severity, as well as its relationship with serum Tau level.

### Limitations

While the current study used state-of-the-art technologies to examine the brain microstructural integrity of adolescent athletes, there were limitations to be noted. A relatively small sample size from a single site, lack of female sports, and imbalance sample size between the football and cross-country groups limit generalizability of the results. We are also aware that the true novelty lies with a longitudinal multimodal relationship, by testing if parameters of neuroimaging and blood biomarkers change over time in relation to head impact exposure. Hirad et al.^47^ recently showed longitudinal agreement between DTI and Tau, but other biomarkers and NODDI were not included. Therefore, this study is an excellent step to encourage interdisciplinary collaborations between neuroimaging and blood biomarker scientists since these fields rarely intersect to delineate subconcussive brain injury. The potential residual neural burden was accounted for by the number of previous concussions and years of tackle football experience. Although these are commonly used variables, there might be an unquantifiable recall bias in self-reporting. A more rigorous approach would be to use head impact data from previous seasons and conduct a medical chart review to validate prior concussion history.

## CONCLUSION

Evidence is beginning to uncover the effects of cumulative concussive and subconcussive head impacts in sports. Neuroimaging and blood biomarker have been two of the most active areas of research in the neurotrauma community. Our data from DTI/NODDI suggest that football players may develop axonal microstructural abnormality. Levels of NDI in the corpus callosum were associated with serum Tau levels, highlighting the vulnerability of the corpus callosum against sport-related head impacts. Despite observing multimodal associations in some brain areas, our study indicates that neuroimaging and blood biomarkers may not strongly correlate to reflect the severity of brain damage. Future study is warranted to determine the longitudinal multimodal relationship in response to repetitive exposure to sport-related head impacts.

## ACKNOWLEDGEMENT

This work was partly supported by the Research Funds from the Indiana University Office of Vice President for Research (to Dr. Kawata), Indiana University Women’s Philanthropy Council (to Dr. Kawata) and Indiana CTSI pilot core fund (to Drs. Kawata and Newman). Sponsors had no role in the design or execution of the study; collection, management, analysis, or interpretation of the data; preparation, review, or approval of the manuscript; or decision to submit the manuscript for publication.

## Author Contribution Statement

KK, JS, JM, AS, SN, HC, ZC, KE conceptualized and designed the study; KK, MH, JS, JM, MN, SN, HC collected the data; KK, SN, ZC, HC, KE, analyzed the data; KK, JS, JM, AS, SN, HC, MN, MH, KK (Kercher) interpret the data; KK, HC, MH, MN drafted the manuscript; JS, JM, AS, SN, KK (Kercher), ZC, KE critically revised the manuscript for important intellectual content; all authors contributed to the final manuscript and interpretation of the final results.

